# *In vivo* visual screen for dopaminergic *Rab ↔ LRRK2-G2019S* interactions in *Drosophila* discriminates *Rab10* from *Rab3*

**DOI:** 10.1101/2020.04.10.035758

**Authors:** Stavroula Petridi, C. Adam Middleton, Chris Ugbode, Alison Fellgett, Laura Covill, Christopher J. H. Elliott

## Abstract

LRRK2 mutations cause Parkinson’s, but the molecular link from increased kinase activity to pathological neurodegeneration remains undetermined. Previous *in vitro* assays indicate that LRRK2 substrates include at least 8 Rab GTPases. We have now examined this hypothesis *in vivo* in a functional, electroretinogram screen, expressing each *Rab* with/without *LRRK2*-*G2019S* in selected *Drosophila* dopaminergic neurons. Our screen discriminated Rab10 from Rab3. The strongest Rab/LRRK2-G2019S interaction is with Rab10; the weakest with Rab3. Rab10 is expressed in a different set of dopaminergic neurons from Rab3. Thus, anatomical and physiological patterns of Rab10 are related. We conclude that Rab10 is a valid substrate of LRRK2 in dopaminergic neurons *in vivo*. We propose that variations in *Rab* expression contribute to differences in the rate of neurodegeneration recorded in different dopaminergic nuclei in Parkinson’s.

## Introduction

Inherited mutations in *LRRK2* (*Leucine-rich-repeat kinase 2*) are a common cause of Parkinson’s. A single amino-acid change, *G2019S*, increases LRRK2 kinase activity (Greggio and Cookson 2009). This mutation results in a toxic cascade that kills *substantia nigra* dopaminergic neurons. However, the main steps in this pathological signalling pathway remain to be determined. Partly this is because LRRK2 is potentially a multi-functional protein, with kinase, GTPase and protein-binding domains. A diverse range of >30 proteins that might be phosphorylated by LRRK2 have been reported, suggesting it is a generalised kinase (Tomkins *et al*. 2018). However, several research teams have recently reported that LRRK2 is a more specific kinase, phosphorylating a range of Rab GTPases (Thirstrup *et al*. 2017; Steger *et al*. 2017; Fan *et al*. 2018; Liu *et al*. 2018; Jeong *et al*. 2018; Kelly *et al*. 2018).

Rabs are a plausible LRRK2 substrate leading to neurodegeneration, as they act as molecular switches interacting with a range of proteins (GEFs, GAPs and GDIs) regulating supply and delivery of cargo to membranes. Indeed many of the 66 Rabs in humans have been linked to neurodegenerative disorders (Kiral *et al*. 2018). Mutations in Rabs 29 and 39 cause Parkinson’s (MacLeod *et al*. 2013; Beilina *et al*. 2014; Wilson *et al*. 2014). Biochemically, at least 8 seem to be directly phosphorylated by LRRK2 [Rabs 3, 5, 8, 10, 12, 29, 35 and 43] (Steger *et al*. 2017). However, it is not clear which of the more than 60 Rabs are actually phosphorylated *in vivo*. In mammals, analysis of the role of the Rabs is complex because individual Rabs may have similar, or even compensatory functions, which may differ by tissue (Chen *et al*. 2012; Kelly *et al*. 2018). The situation is simpler in the fly, because there are fewer Rabs - only 23 mammalian orthologs. Here, we use a *Drosophila* screen to assess the link from LRRK2 to Rabs *in vivo* using the *Tyrosine Hydroxylase* (*TH*) GAL4 to achieve dopamine specific expression. *UAS-LRRK2-G2019S* (Liu *et al*. 2008) was driven with and without each *Rab* gene (Zhang *et al*. 2006).

We measured a visual phenotype using the SSVEP (Steady State Visual Evoked Potential). Although the outer structure of the eye differs markedly between flies and mammals, the retinal circuitry is highly similar (Cajal and Sanchez 1915; Sanes and Zipursky 2010) – importantly both contain dopaminergic neurons. In the human, the retinal dopaminergic neurons die in Parkinson’s (Harnois and Di Paolo 1990), while in the *TH>G2019S* model of Parkinson’s, the retina has visual deficits, including neurodegeneration (Hindle *et al*. 2013; Afsari *et al*. 2014; West *et al*. 2015a). We can now use the ability of the SSVEP assay to separate and quantify the response of the photoreceptors and lamina neurons to go beyond measuring neurodegeneration, but to test for a synergistic interaction of a Parkinson’s related gene with potential substrates. Notably, we can do this *in vivo* in young flies before degeneration has set in.

We determined that, *in vivo*, Rab10 has the strongest synergy with LRRK2-G2019S, Rab3 the weakest. We validated the physiological results by showing differences in the expression of *Rab10* and *Rab3* in visual dopaminergic interneurons.

## Materials and methods

### Flies

(*Drosophila melanogaster*) were raised and manipulated according to standard fly techniques. Fly stocks are listed in Table 1. Crosses were raised at 25 °C on a 12:12 light-dark cycle. On the day of emergence, female flies were placed in the dark at 29 °C.

### Screen design

Virgins from the *TH*-GAL4, or from a *TH*-GAL4::UAS-*LRRK2*-*G2019S* (*THG2*) recombinant were crossed with males carrying UAS-*Rab*, for each of the *Rabs* that are homologous to those of mammals.

The principle of the SSVEP screen is shown in Fig. 1. The visual response of flies stimulated with a flickering blue light was recorded. Young, 4-12 hour old, PD-mimic flies show visual hyperexcitability, particularly in the lamina neurons (Afsari *et al*. 2014; Himmelberg *et al*. 2017). This includes the *THG2* flies. As they age, the visual response gets weaker and vanishes by 28 days. We therefore chose to test flies aged for 24-36 hours (1 day) or 1 week - between the time at which *G2019S* expression results in hyperexcitability and the time at which degeneration is first noted. At these time points, the mean visual response of dark-reared *THG2* flies was similar to the *TH/+* controls.

**Fig. 1.**
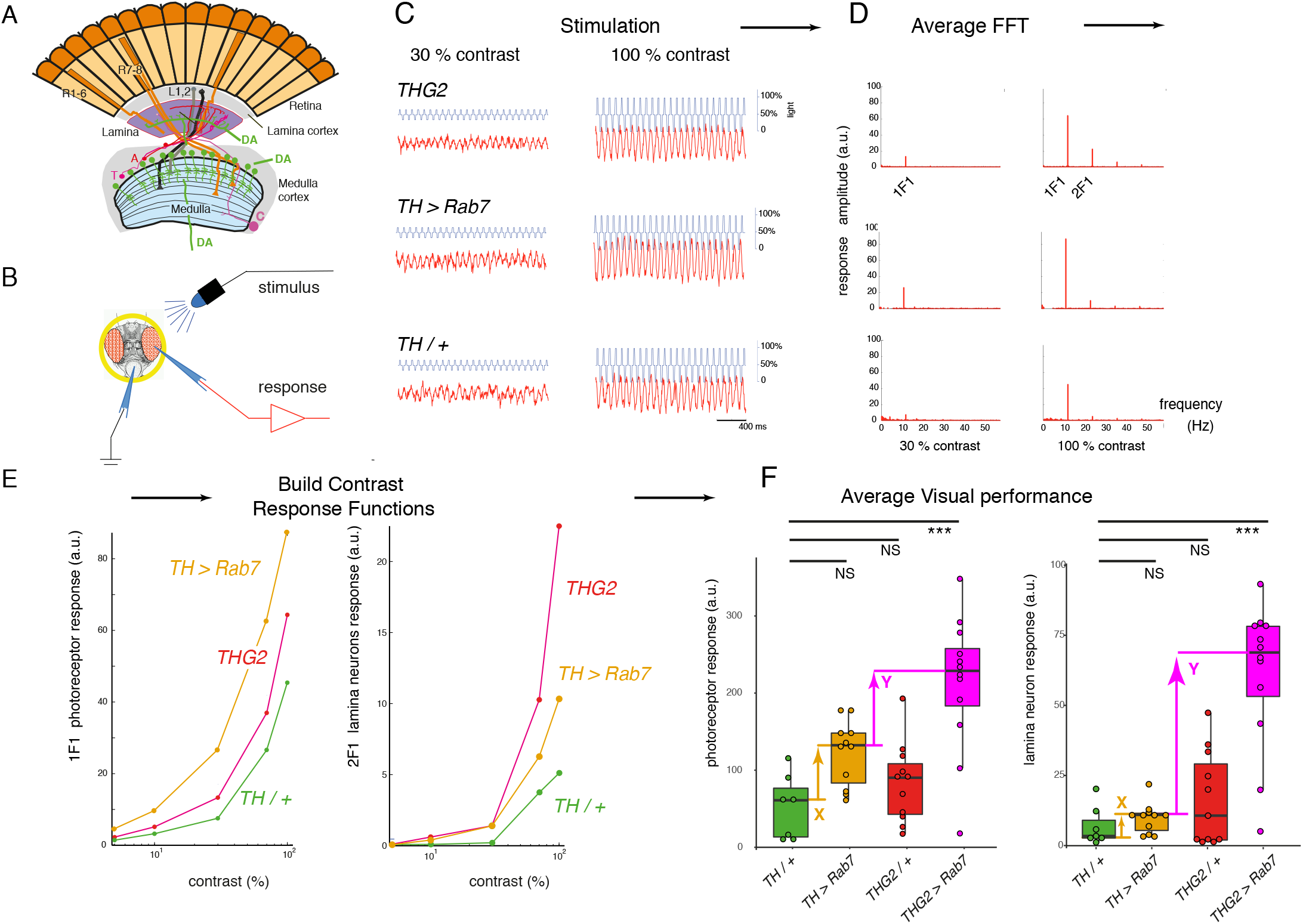
SSVEP (Steady State Visual Evoked Potential Analysis) measures the contrast response function of the insect eye. A. The fly eye consists of ~800 ommatidia, each containing 8 photoreceptors. Their axons project to the lamina and medulla, where they synapse with the second- and third order neurons (lamina and medulla neurons). The medulla contains intrinsic dopaminergic neurons (MC, also called Mi15 neurons (Davis *et al*. 2020)), while some dopaminergic neurons from the CNS project to the lamina. B. Recording the fly visual response: A fly, restrained in a pipette tip, is illuminated with blue light from a LED, and the voltage across the eye is amplified and recorded. C. Repetitive stimuli given to the fly about a fixed mean light level evoke a contrast response increasing with the peak-peak excursion of the stimulus waveform. D. The response to a series of identical stimuli is analysed by the Fast Fourier Transform, and averaged. This shows a response at the stimulus frequency (1F1) and additional components at multiples of the input, notably twice the input frequency (2F1). Genetic dissection shows that the 1F1 component is mostly generated by the photoreceptors and the 2F1 by the lamina neurons (Afsari *et al*. 2014; Nippe *et al*. 2017). E. Plotting the amplitude of the 1F1 and 2F1 components against the stimulus contrast generates a CRF (Contrast Response Function), which differs from fly to fly. F. The averaged maximum CRF is dependent on genotype, with *THG2* (flies expressing *LRRK2-G2019S* in their dopaminergic neurons under the *Tyrosine Hydroxylase*-GAL4, *TH*) and *TH* > *Rab7* both showing a similar mean response to control flies (*TH/+*). However, flies expressing both *G2019S* and *Rab7* in their dopaminergic neurons (*THG2* > *Rab7*) have a larger mean response than any other genotype, indicating synergy. The differences marked X (between the mean *TH/+* and *TH* > *Rab7*) and Y (between the mean *TH* > *Rab7* and *THG2* > *Rab7*) are used as the X and Y axes of Fig. 2A. Box-plot representing median and interquartile range. Exact genotypes and sample sizes: *TH/+, TH/w^1118^*, N= 7; *THG2, TH::G2019S/ w^1118^*, N=11; *TH* > *Rab7*, N=11; *THG2* > *Rab7*, N=12.

### Sample test for synergy

We test for an interaction between *Rab7* and *G2019S* in dopaminergic neurons as follows: we compare flies expressing both *Rab7* and *G2019S* transgenes (*THG2* > *Rab7*) with flies expressing just one transgene (*TH* > *Rab7* or *THG2*) and control flies with no transgene expression (*TH/+*) (Fig. 1F). The average visual response of *TH* > *Rab7* and *THG2* flies is very similar to the control flies – there is no mean difference for either the photoreceptors or lamina neurons. We do note that the *THG2* flies have a larger variability than the *TH/+* flies, particularly in the lamina neurons (Fig. 1F). However, in flies with dopaminergic expression of both *G2019S* and *Rab7*, the photoreceptor and lamina neuron responses were much increased (4.1x and 8.8x, both P < 0.001). This demonstrates that dopaminergic neurons with both *Rab7* and *G2019S* have a synergistic hyperexcitable visual phenotype.

### SSVEP preparation

At the required age, flies were prepared for SSVEP measurements using a pooter and nail polish to secure them in the cut-off tip of a pipette tip, without anaesthesia (Fig. 1B). Each fly was presented 5 times with a set of 9 flickering stimuli. In each stimulus, the average light intensity was the same, but the amplitude of the flicker was adjusted from 10 to 100%, giving a range of contrasts. Sample stimuli are shown in Fig. 1C. Offline, the Fast-Fourier Transform was applied to the responses, to separate the first harmonic (1F1), due to the photoreceptors from the second harmonic (2F1), due to the lamina neurons (Fig. 1D). Other harmonics present in the data were not analysed. For these first two harmonics, we plotted the contrast response function for each fly (Fig. 1E) and determined the best response of that fly. This allowed us to determine the average visual performance for each cross (Fig. 1F). This data pipeline is the same as that devised by Afsari *et al*. (2014), but using an Arduino Due to generate the stimuli and record the responses instead of a PC. Data were analysed in Matlab, Excel and R. Full code at https://github.com/wadelab/flyCode.

### Immunocytochemistry

was performed as described recently (Cording *et al*. 2017). Tyrosine hydroxylase was detected with Mouse anti TH Immunostar (22941, 1:1000). Fluorescent reporters (nRFP, eIf-GFP) were expressed in dopaminergic neurons using the *TH*-GAL4. Images were prepared for publication using ImageJ; original images are available on request.

### Western blots for

EYFP, encoded in each Rab transgene were made from non-boiled fly head lysates, run on Novex pre-cast mini gels (NuPAGE 4-12% Bis-Tris Gels, NP0322BOX, Thermo Scientific) in 1 x MOPS buffer and transferred onto PVDF membranes using a Hoefer wet transfer tank (TE22) at 100V for 1 hr. Membranes were probed with Guinea pig anti-GFP (Synaptic Systems, 1:1000). For detection of LRRK2 protein, boiled lysates were run on 4-20% Mini-PROTEAN TGX Precast gradient gels and transferred using the same method. Membranes were probed with anti-LRRK2 (Neuromab, clone N241A/34, 1:1000). α-drosophila synaptotagmin was used as a loading control (West *et al*. 2015b). Densitometric analysis was carried out using ImageJ.

### Statistics

were calculated in R, with the mean ± SE reported by error bars or median ± interquartile range in box plots. Post-hoc tests were calculated for ANOVA using the Dunnett test.

### Data Availability Statement

Data tables (Excel sheets) and R code are open access on GitHub: https://github.com/wadelab/flyCode/tree/master/analyzeData/fly_arduino/G3. Raw images and SSVEP traces are available on request. No new reagents are described.

## Results

### A visual expression screen identifies that Rab10 has the strongest genetic interaction with LRRK2-G2019S; Rab3 the weakest

In order to identify the Rabs which show a strong synergy with *LRRK2*-*G2019S* we compared the increase in visual response due to expression of the *Rab* by itself (X in Fig. 1F, X-axis in Fig. 2A) against the further increase in visual response when both *Rab* and *G2019S* are expressed (Y in Fig. 1F, Y-axis in Fig. 2A). This plot places the Rabs along a spectrum, from those that interact synergistically with *G2019S* (top left) to those with a little or no interaction (bottom right). Thus, for some Rabs (10, 14, 27, 26) expression of both *G2019S* and the *Rab* in dopaminergic neurons leads to a big increase in the lamina neuron response. Interestingly, these Rabs have little effect when expressed alone. The converse is also true: for the *Rabs* with the biggest effect (3, 32, 1), adding *G2019S* has no further effect. This is true for both components of the SSVEP signal – that from the lamina neurons is higher, but parallel to the photoreceptor signal.

**Fig. 2.**
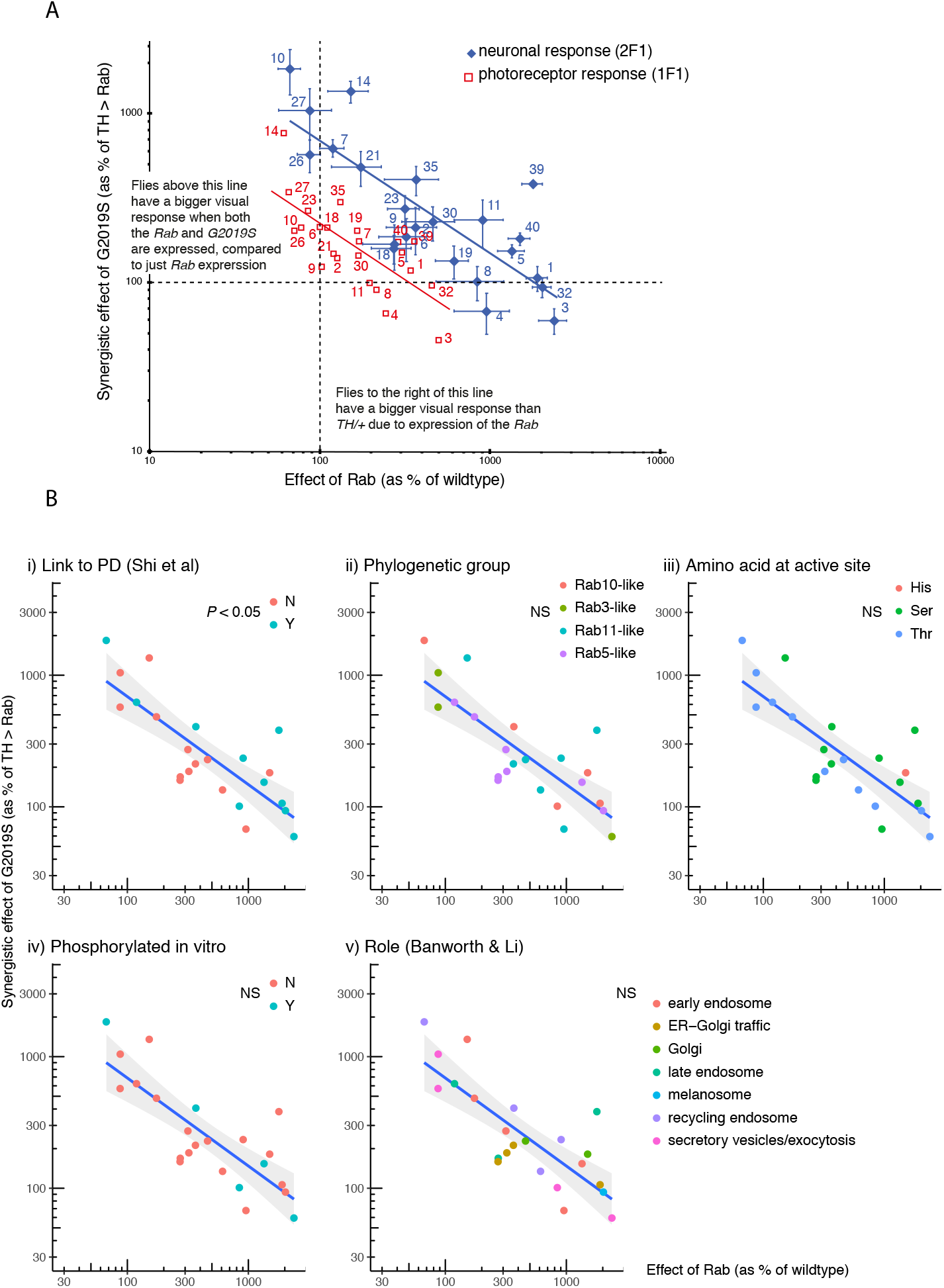
Expression screen highlights *Rab10* with the strongest synergy with *LRRK2-G2019S*, and *Rab3* as the weakest. Each *Rab* was expressed in dopaminergic neurons (using *TH*-GAL4) by itself (*TH* > *Rab*), or along with *G2019S* (*THG2* > *Rab*) and the visual response measured after 24-36 hours (labelled 1 day) or 7 days in the dark. A. **Rab10 has the strongest synergy with *G2019S***. Relationship of *Rab* and *G2019S* showing their inverse relationship. Rabs (3, 32, 1) which have a big effect on vision when expressed on their own have little further consequence when *G2019S* is also expressed; but other Rabs (10, 27, 14, 26) which have little visual impact on their own have a strong synergy with *G2019S*. B. **An established role in Parkinson’s is the only factor that influences the inverse relationship between *TH* > *Rab* and *THG2* > *Rab***. The LRRK2 **↔** Rab data in Fig. 2A are replotted here to test if it is affected by factors that have been proposed to influence LRRK2 **↔** Rab interactions. (i). Rabs previously linked to Parkinson’s (Shi *et al*. 2017) have a stronger Rab **↔** G2019S response than those which do not influence Parkinson’s, since a higher proportion of the magenta points lie above the line (Fisher’s exact test, P = 0.036). B(ii). Sensitivity is not linked to the phylogenetic grouping of the fly Rabs (Zhang *et al*. 2006). B(iii). Rabs usually have a serine (Ser) or threonine (Thr) where they could be phosphorylated by LRRK2, though Rab40 has a histidine (His) (Zhang *et al*. 2006). Although a preference for LRRK2 to phosphorylate Rabs with a threonine was suggested by *in vitro* assays (Steger *et al*. 2016), *in vivo* this is not detected. B(iv). Some Rabs are phosphorylated by LRRK2 *in vitro* (Steger *et al*. 2017), but these Rabs are not more sensitive to *G2019S in vivo*. B(v). The proposed main functional role of the Rab (Banworth and Li 2017) does not affect the regression. Total flies: 1119, at least 9 for each data point. Bars represent SE.

We wanted to examine which factors controlled this synergy. A number of hypotheses have been put forward in the LRRK2/Rab literature. First, Rabs previously linked to Parkinson’s (Shi *et al*. 2017), either through population studies or through potential actions with Parkinson’s-related genes, generally have a stronger response to *G2019S* than others (Fig. 2Bi). Indeed, the Rab furthest above the regression line is one that causes Parkinson’s, *Rab39* (Wilson *et al*. 2014). Next, we tested if Rabs with a high degree of phylogenetic similarity clustered systematically, but did not find any difference (Fig. 2Bii). Then we examined where, on our spectrum, the Rabs phosphorylated *in vitro* [3, 5, 8, 10, 12, 29, 35 and 43] might lie. There is no close homolog for Rabs 12, 29 or 43. Rabs 3, 5 and 8 are on the right of the spectrum, Rab35 in the middle and Rab10 on the top-left, so no clear pattern emerges. The *in vitro* data suggest that LRRK2-G2019S preferentially phosphorylates Rabs with Thr rather than Ser at the active site (Steger *et al*. 2016), but this is not evident from the spectrum. Neither *in vitro* evidence for phosphorylation of the Rab by LRRK2, nor the amino-acid at the active site affects the regression (Fig. 2Biii,iv). We also noted that, in cell based assays, LRRK/Rab interactions have been noted at mitochondria (Wauters *et al*. 2019), lysosomes (Eguchi *et al*. 2018) or Golgi (Liu *et al*. 2018). Thus, we tested if the ‘main’ organelle associated with the Rab (Banworth and Li 2017) affected the position of a Rab on the spectrum, but found no sign that this affected the regression (Fig. 2Bv). However, the Rabs placed in the middle of the spectrum [2, 6, 9, 18] are linked to traffic in the Golgi or ER-Golgi.

Thus, the relationship between visual impact of *Rab* and impact of *Rab* + *G2019S* identifies 10, 14, 27 as having the strongest synergy. This holds for the responses of both photoreceptors and lamina neurons, with the same Rabs found clustered at each end of the spectrum.

### The Rab10/G2019S interaction enhances neuronal signalling

Normally, flies with more excitable photoreceptors activate the lamina neurons more strongly, though there is some adaptation. The SSVEP response can be decomposed into two components – 1F1 and 2F1, corresponding to activity in the photoreceptors and lamina neurons respectively. This allows us to test the physiological relationship between the photoreceptors and lamina neurons, and to see if any Rab disrupts the retinal neuronal circuitry. Generally, as the photoreceptor response increases, so does the lamina neuron response (Fig. 3). This relationship is remarkably similar in young (day 1) and older (day 7) flies. However, there is a one marked outlier, *Rab10*, where the lamina neuron response at day 1 is ~5 times the value expected from the regression, and at day 7 is substantially below the line. Thus, in young *THG2* > *Rab10* flies there is much greater neuronal activity than expected, but in 1-week old *THG2* > *Rab10* flies we observe reduced activity, suggesting neurodegeneration has begun. Young and old Rab3 flies lie on the regression, close to the origin, very different from Rab10.

**Fig. 3.**
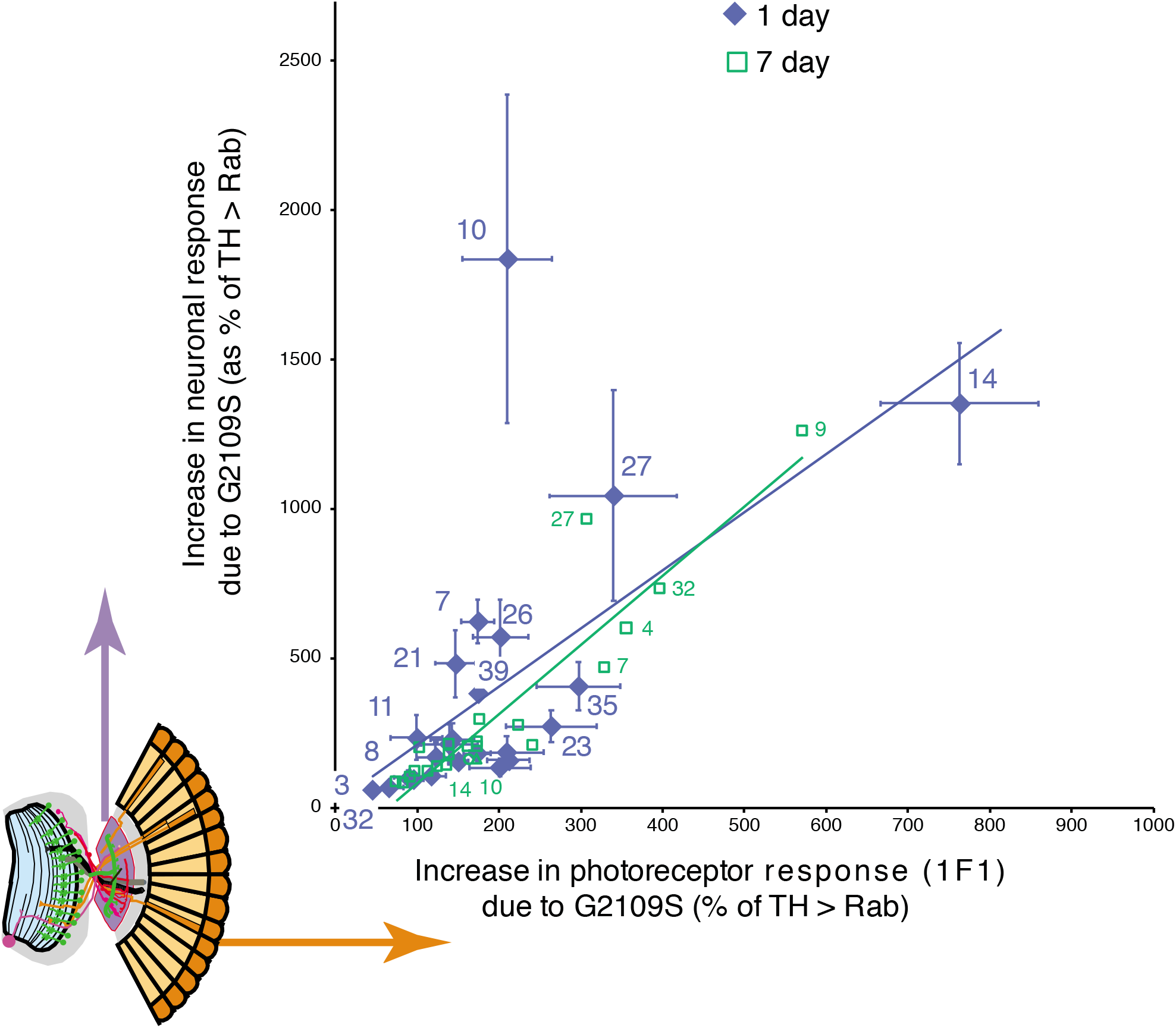
Standout role of *Rab10* with *G2019S* in neuronal signaling. The SSVEP response is split into two components, representing the photoreceptors and lamina neurons (inset orange and purple). For each Rab, the increase in lamina neuronal signaling due to *G2019S* is plotted as a function of the photoreceptor signal. The increase in lamina neuron response is highly correlated with the response of the photoreceptors, with one outlying exception, *Rab10* at 1 day. Total flies: 1119, at least 9 for each data point. Bars represent SE.

Thus our screen highlights a major difference between two of the Rabs that are phosphorylated in vitro: *Rab3* expression in dopaminergic neurons has a big increase in visual sensitivity, but no further effect when *G2019S* is added, whereas Rab10 expression has little effect by itself, but a massive effect in young flies with *G2019S*.

### Why might G2019S interact so strongly with Rab10 but have no effect on Rab3?

As LRRK2 is a human protein, and the Rabs we expressed were native *Drosophila* proteins, one possibility is that the fly and human Rabs are sufficiently different that LRRK2-G2019S can phosphorylate fly Rab10 but not fly Rab3. This seems very unlikely as the hRab3 / dRab3 and dRab10 / hRab10 sequences are very similar, indeed they are identical over the GTP binding domain and LRRK2 phosphorylation sites (Fig. 4). A second explanation for the difference between Rab10 and Rab3 is that the *Rab* and/or *G2019S* is not expressed to the same extent. Western Blots of *THG2* > *Rabs 3, 39* or *10* were probed for LRRK2, and compared with *THG2*. Essentially the same level of protein was measured (Fig. 5A). This is not unexpected, as each cross contains the same GAL4 and UAS-*LRRK2*-*G2019S* constructs. A second set of blots were probed for EYFP, as each of the UAS-*Rab* lines carries an EYFP fusion. This showed that the levels of Rab10 and Rab39 were similar, though Rab3 was less at about 50% (Fig. 5B). The reduced level of Rab3 may arise from the different insertion site, or from a more rapid breakdown during synaptic signalling. The differences in level of Rab proteins are not sufficient to explain the physiological differences.

**Fig. 4.**
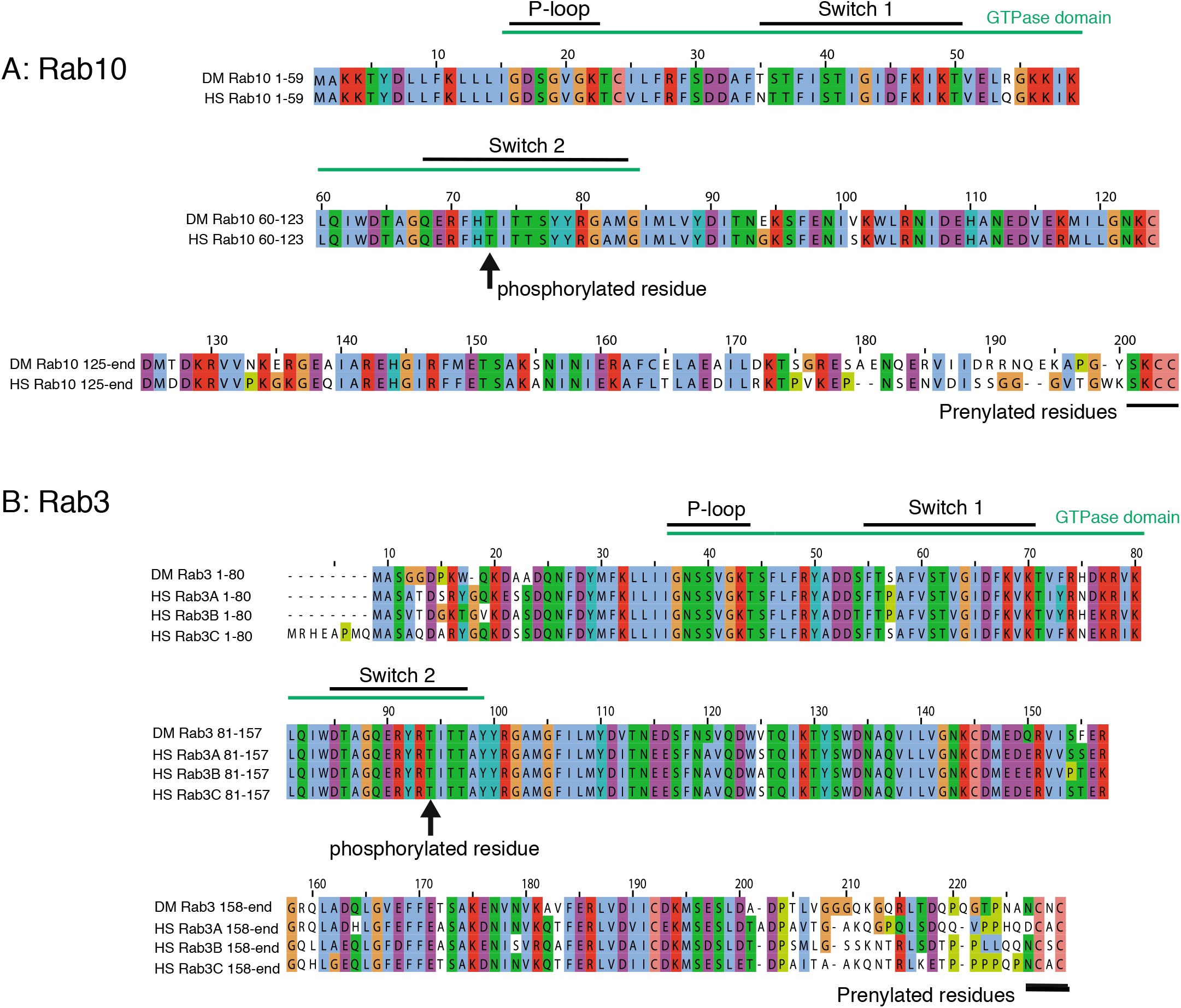
High sequence homology between *Drosophila* and human Rabs. Comparison of fly and human Rab10 (A) and Rab3 (B) showing conservation in the GTPase domain and prenylated region. Also shown is the region which is phosphorylated *in vitro* by LRRK2, again highly conserved.

**Fig. 5.**
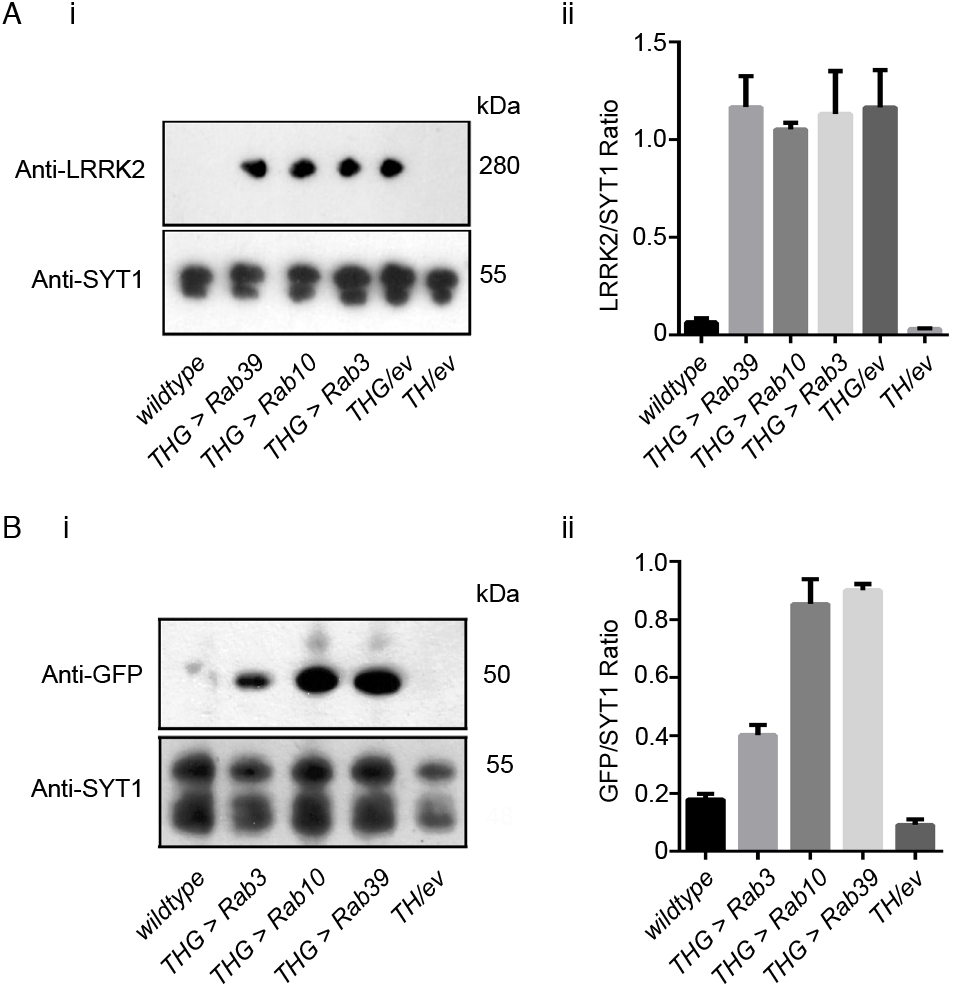
Similar Expression of LRRK2-G2019S and Rab-GFP in dopaminergic neurons. A. Co-expression of a *Rab-GFP* transgene does not affect the levels of *LRRK2-G2019S*. (i) Sample blot, (ii) Quantification of 3 replicates. B. Similar levels of Rab10 and Rab39, and less Rab3 when driven with *LRRK2-G2019S*. (i) Sample blot, (ii) Quantification of 3 replicates. wild-type is *CS/w*^−^, *TH/+* is *TH/empty vector*.

We therefore wondered if the stronger synergy between G2019S and Rab10, compared with Rab3, might result from a difference in the anatomical distribution of the Rabs (along with their GEFS, GAPs and effectors) among fly dopaminergic neurons.

### Rab10 and Rab3 are found in different dopaminergic neurons

The fly visual system is innervated by three kinds of dopaminergic neurons (Hindle *et al*. 2013), the MC neurons in the medulla, and two type of PPL neurons, which innervate either lobula or lamina respectively. These, and the other clusters of dopaminergic neurons, are reliably marked by α-TH antibody, which binds in the cytoplasm.

To examine the overall distribution of Rab10, we used *Rab10-GAL4* (Chan *et al*. 2011) to express either a RFP which strongly localises to the nucleus, or a GFP with mainly nuclear localisation. These fluorescent constructs have two advantages: (i) they provide a reduced background compared with membrane localised reporters, and (ii) the nuclear fluorescence is contained within the cytoplasmic signal from α-TH, reducing the problems of determining co-localisation.

Only a small proportion of CNS neurons are Rab10 positive (Fig. 6). We find that some (by no means all) dopaminergic neurons are Rab10 positive (Fig. 6A,B). Even within a cluster, we only detect Rab10 in some neurons; in other neurons in the same cluster Rab10 is undetectable (e.g. PAL, PPL2ab, PPM3 and PAM). The individually identifiable neurons (TH-VUM, TH_AUM, the DADN pair, and T1 pair) were consistently clearly marked. However, in two clusters we saw no evidence for *Rab10* driven fluorescence (PPL2c and PPM1/2).

**Fig. 6.**
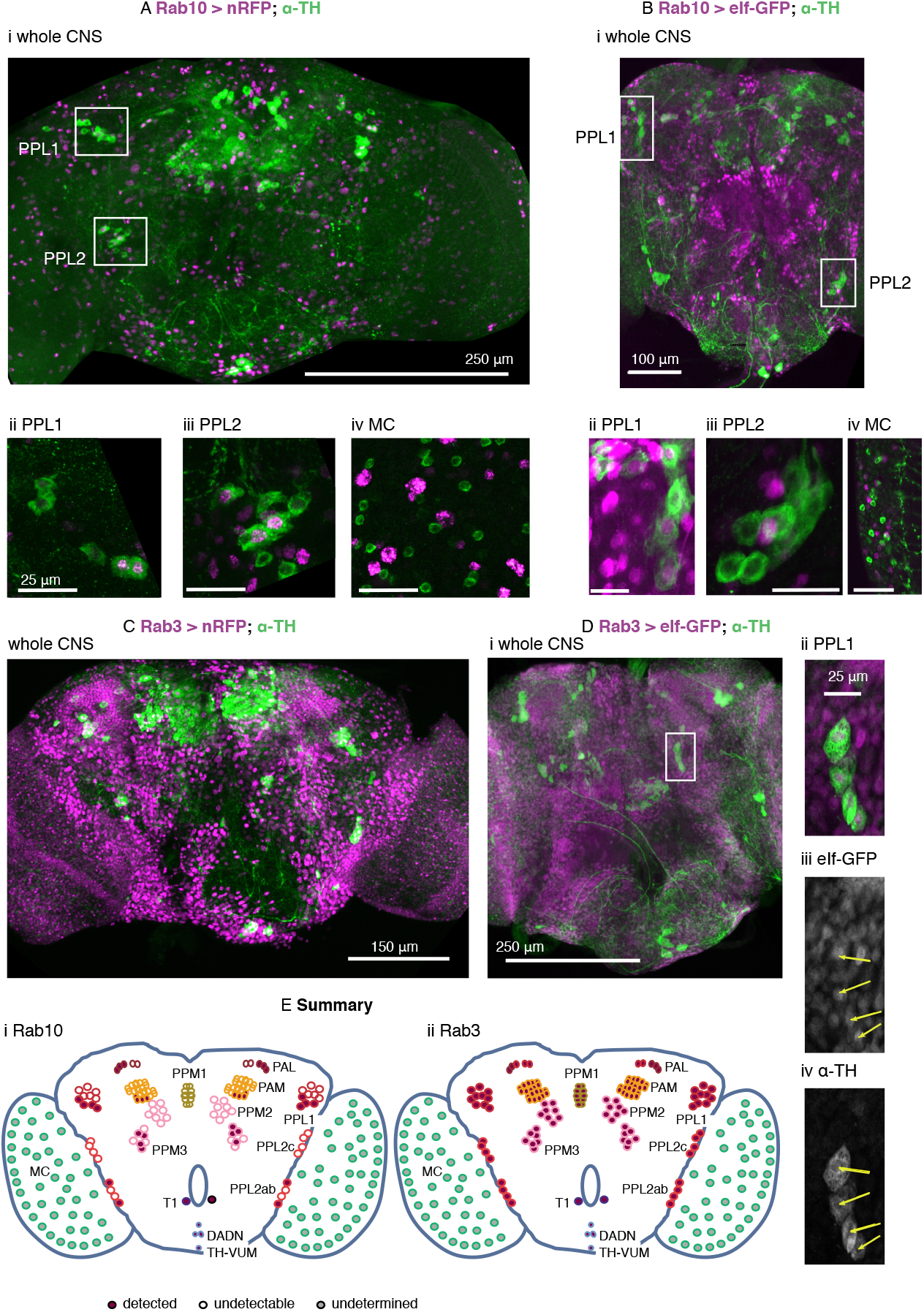
Rab10 and Rab3 are located in different subsets of the dopaminergic neurons. A, B. Rab10 is detected in some of the dopaminergic neurons that control vision (PPL1, Aii, Bii; PPL2 Aiii, Biii). Not all dopaminergic neurons, identified by a cytosolic α-Tyrosine Hydroxylase antibody (α-TH, green), are indicated by *Rab10*-GAL4 expression of a strong nuclear RFP or the mainly nuclear eIf-GFP (magenta). The dopaminergic MC neurons in the visual lobes do not stain well with fluorescent reporters (Nassel and Elekes 1992; Hindle *et al*. 2013) and we could not detect *Rab10*-driven fluorescence (MC, Aiv, Biv, marked with grey in E). C, D. **Rab 3 is present in all dopaminergic neurons**. *Rab3*-GAL4 driven nuclear RFP or eIf-GFP (magenta) marks most neurons, including nearly all that are dopaminergic (green). The PPL neurons not marked by *Rab10* expression are included (Dii-iv). E. Summary of the expression pattern of (i) *Rab10* and (ii) *Rab3*. The MC neurons in the optic lobe (Nassel *et al*. 1988) are also called Mi15 neurons (Davis *et al*. 2020). Ai, Bi, Ci and Di: projection of confocal stacks through the whole CNS; Aii, Aiii, Bii, Biii, Dii-iv projections of confocal stacks through the cell groups, approximately marked in the whole CNS image; Aiv and Biv sections from a separate preparation to Ai and Bi. Data representative of at least nine brains (from at least 3 crosses), 3-7 days old. The *Rab3* > *nRFP* flies were raised at 18 °C to improve viability. Exact genotypes: +; *RedStinger4 nRFP/+; Rab10 Gal4/+;* or *+; RedStinger4 nRFP/+; Rab3 Gal4/+;* or +; *eIf-4A3-GFP/+; Rab10 Gal4/+;* or +; *eIf-4A3-GFP/+; Rab3 Gal4/+*;

When we used *Rab3*-GAL4 to drive the same RFP/GFP almost all the neurons were marked (Fig. 6 C, Di). This includes the majority of the dopaminergic neurons, including all the PPL1 (Fig. 6 Dii-iv) and PPL2 neurons.

The MC neurons in the optic lobes were not marked in either the Rab10 or Rab3 experiments (Fig. 6 Aiv, Biv), though other Rab10 / Rab3 positive neurons are present nearby. Since the MC neurons do not generally stain well with GFP (Nassel and Elekes 1992; Hindle *et al*. 2013)), we tested if the MC neurons were detected with *TH*-GAL4 > *nRFP*. This marked all the neurons highlighted by α-TH, except the MC neurons. The MC neurons do express

TH, along with other genes linked to dopamine - *ddc* (*dopa decarboxylase*), *Vmat* (*vesicular monoamine transporter*) and *DAT* (*dopamine transporter*) (Davis *et al*. 2020) so are genuinely dopaminergic. The MC neurons are one of three kinds of Medulla intrinsic neurons that express *Rab10* at high levels, while all the optic lobe neurons (including MC) have high expression of Rab3 (Davis et al, 2020, extended data at http://www.opticlobe.com/).

Thus, we conclude that some of the dopaminergic neurons in the visual system are Rab10 positive. These are some of the PPL cluster that innervate the lobula or project to the lamina, and the MC neurons in the medulla. All dopaminergic neurons are Rab3 positive.

### Differences in the loss of dopaminergic neurons between neuronal clusters

*Drosophila* models of Parkinson’s have consistently shown loss of dopaminergic neurons with age when *LRRK2, α-synuclein* or *parkin* were manipulated. For *LRRK2*, most of the published information is for the Parkinson’s-causative mutations *G2019S* or *I2020T*, driven by *DDC*-GAL4. This expresses in the dopaminergic and some serotonergic neurons. By 6-7 weeks (about two-thirds of the fly lifespan), about 25-50% of the dopaminergic neurons have been lost. For each cluster, there is quite a spread of the data (Fig. 7), which is most likely due to differences in the food used to feed the flies or the genetic background (Lavoy *et al*. 2018; Chittoor-Vinod *et al*. 2020). However, overall, the PAL cluster is much less susceptible to cell loss than the PPL1, PPL2, PPM1/2 or PPM3 clusters.

**Fig. 7.**
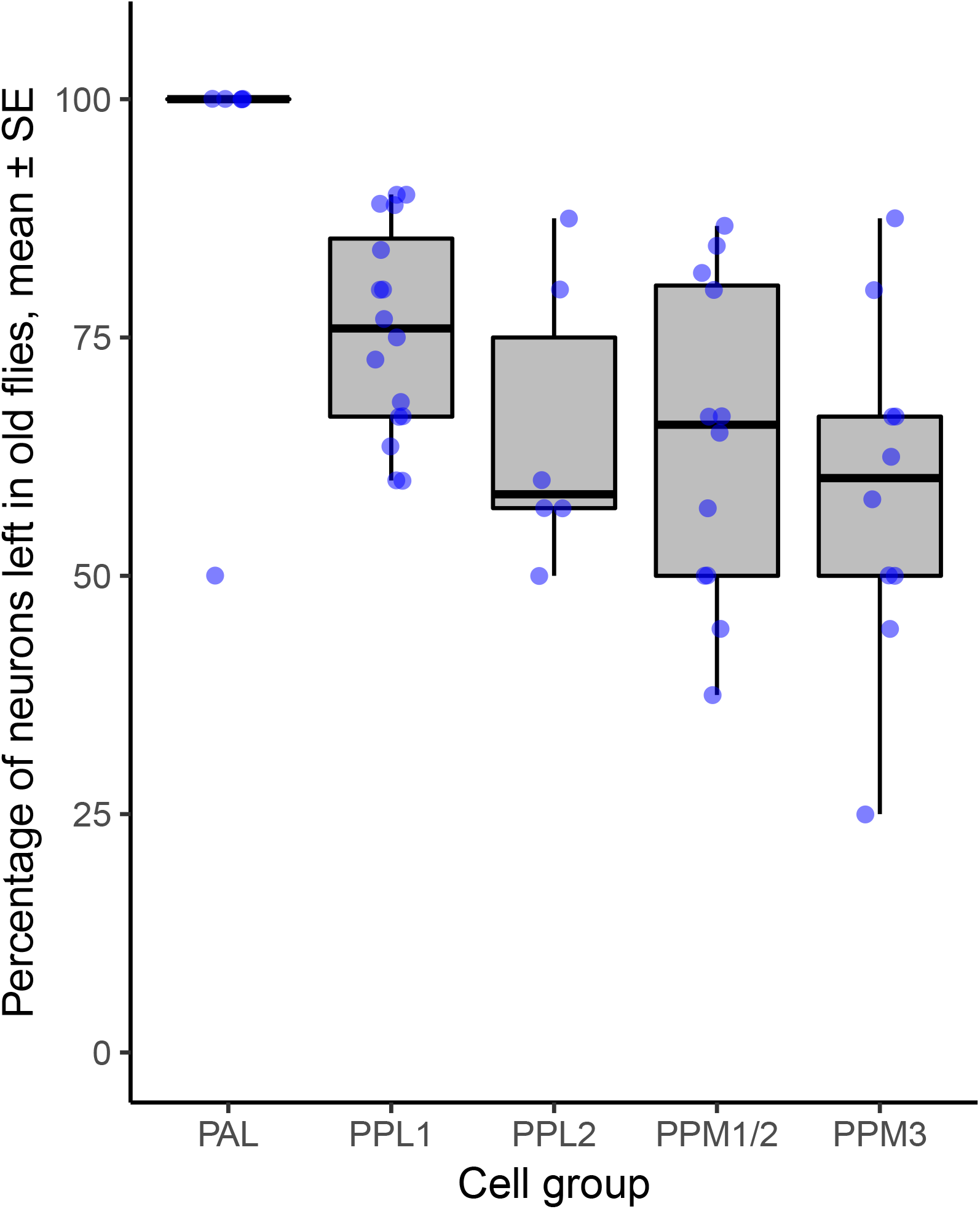
Differences in neuron survival in dopaminergic clusters when an increased kinase mutation (*G2019S* or *I2020T*) is expressed with *DDC*-GAL4 (which expresses in dopaminergic and serotonergic neurons) (ANOVA, 4,45 df, *P*<0.002). Data collected from (Liu *et al*. 2008; Ng *et al*. 2009; Xiong *et al*. 2012; Angeles *et al*. 2014, 2016; Martin *et al*. 2014; Nucifora *et al*. 2016; Lin *et al*. 2016; Sun *et al*. 2016; Basil *et al*. 2017; Marcogliese *et al*. 2017; Yang *et al*. 2018; Lavoy *et al*. 2018; Sim *et al*. 2019; Maksoud *et al*. 2019; Chittoor-Vinod *et al*. 2020). Differences in the extent of degeneration within a neuronal cluster may be partially explained by differences in the composition of the fly food (Chittoor-Vinod *et al*. 2020).

## Discussion

### Rab10 shows a strong synergy with LRRK2-G2019S

The key observation from the screen was that two of the Rabs suggested to be substrates of LRRK2 *in vitro* behave quite differently *in vivo*, in a physiological response to expression in dopaminergic neurons. Rab10 shows a strong synergy with *G2019S;* Rab3 none. The existence of (*Drosophila*) Rab10 in the tyrosine hydroxylase positive neurons controlling vision (MC and PPL2ab neurons) argues that LRRK2 might indeed phosphorylate dRab10 directly. Thus our *in vivo* results both support the *in vitro* (biochemical and cell culture) data in which LRRK2 directly phosphorylates hRab10 (Thirstrup *et al*. 2017; Steger *et al*. 2017; Fan *et al*. 2018; Liu *et al*. 2018; Jeong *et al*. 2018; Kelly *et al*. 2018). It also implies that the MC / PPL2ab cells contain Rab10 effectors which interact with phospho-dRab10. The results of this will include changes to the cellular homeostasis and physiological responsiveness of dopaminergic neurons. One possible physiological outcome is that Rab10 phosphorylation reduces retinal dopamine release onto the photoreceptors. This will increase the amplitude and speed of the photoreceptor response (Chyb *et al*. 1999). Dopamine may also affect the lamina neurons, and third order MC cells, but it remains to be determined if they have dopamine receptors. It is also possible that p-Rab10 modulates the release of co-transmitters or growth factors from dopaminergic neurons.

A unique feature of the screen is that when *G2019S* and *Rab10* are expressed together the lamina neuron response is much bigger than that predicted from the photoreceptor response. This might arise from the unusual double role of Rab10 – in both exo- and in endo-cytosis (Larance *et al*. 2005; Glodowski *et al*. 2007; Chua and Tang 2018). The best defined role of Rab10 in exocytosis is in adipocytes, as part of the insulin-stimulated release of GLUT4 vesicles, linked to AS/160 (see for review (Jaldin-Fincati *et al*. 2017)). In endocytosis, the effects of Rab10 are mediated through a different pathway, including the EHBP1-EHD2 complex. In the follicle cells of *Drosophila*, *ehbp1* expression and knockdown phenocopy Rab10 manipulations (Isabella and Horne-Badovinac 2016), while EHBP1 was also identified by a systematic proteomic analysis as indirectly phosphorylated by LRRK2 in HEK293 cells (Steger *et al*. 2017) and a lysosomal assay (Eguchi *et al*. 2018). The phosphorylation of Rab10 by LRRK2 may switch its effector, and so activate a different pathway.

### A spectrum of Rab ↔ G2019S interactions in vision

Our screen placed the Rabs along a spectrum, ranging from those with a strong synergy with *G2019S* to those which had a strong effect when expressed by themselves.

Among the Rabs which show little synergy with *G2019S* but have strong visual effect are 1, 3, 5, 6 and 11. Two of these Rabs [3,5] are phosphorylated by LRRK2 *in vitro* (Steger *et al*. 2017), but neither synergise with LRRK2-G2019S in the visual assay. Our data suggest Rab3 is not a major substrate of *LRRK2-G2019S* in these dopaminergic neurons, possibly because Rab3 is located synaptically. This may be far from LRRK2 at the trans-Golgi network (Liu *et al*. 2018). The difference between Rab3 and 10 (at opposite ends of our spectrum) is notable because *in vitro* mammalian cell assays have highlighted similar roles of Rabs 3 and 10 in lysosome exocytosis, (Encarnação *et al*. 2016; Vieira 2018).

Rabs 10, 14 and 27 have the strongest synergy with *G2019S*, though by themselves they have little effect on visual sensitivity. Like Rab10, Rabs 14 and 27 have defined roles in exocytosis (Larance *et al*. 2005; Ostrowski *et al*. 2010).

Some Rabs are in the middle of the spectrum [2, 6, 9, 18], with a 2-3 fold increase in visual response when the *Rab* is expressed alone, and a further 2-3 fold increase when both *Rab* and *G2019S* are expressed. These Rabs have been linked to the Golgi, or to Golgi-ER traffic (Banworth and Li 2017). Thus a cellular phenotype parallels the physiological response.

Our observation that every Rab seems to have some effect on dopaminergic signalling in the visual system goes some way to explain why many studies of individual Rabs have demonstrated effects with LRRK2; Rab3a (Islam *et al*. 2016); Rab5 (Shin *et al*. 2008); Rab7 (Dodson *et al*. 2012); Rab29 (Beilina *et al*. 2014). Although cellular studies support binding of Rab29 to LRRK2 (Purlyte *et al*. 2018), the closest fly homolog (Rab32) shows little synergy with *G2019S* in our screen.

The availability of Rab transgenic flies facilitates screening in *Drosophila*. Screens have identified key roles for Rab2 in muscle T-tubule development (Fujita *et al*. 2017); Rabs 2, 7, 19 in loss of huntingtin (White *et al*. 2015), 1, 5, 7, 11 and 35 in the *Drosophila* renal system (Fu *et al*. 2017), Rab32 in lipid storage (Wang *et al*. 2012) and Rab39 in tracheal formation (Caviglia *et al*. 2016). The varied outcomes of these screens indicate the validity of the LRRK2-G2019S ↔ Rab10 relationship reported here.

### Each dopaminergic neuron has its own palette of Rab expression

Finally, we note that not all dopaminergic neurons are equally susceptible in Parkinson’s. A long-standing observation is that the dopaminergic neurons in the VTA (ventral tegmental area) do not degenerate in the same way as those in the *substantia nigra*. More particularly, even within the *substantia nigra* there is a range of outcomes, with dopaminergic neurons in the *pars compacta* dying more than those in the dorsal and lateral zones (Damier *et al*. 1999). The same is true for the fly brain: the neurons in the PPM clusters degenerate more than the PAL (though no data are available for the visual MC neurons). If anything, our data suggest the clusters with less Rab10 have more neurodegeneration. Previously, faster neurodegeneration has been ascribed to increased cytosolic dopamine levels (Burbulla *et al*. 2017), to intracellular effects of glutamate (Steinkellner *et al*. 2018), to increased calcium influx (Guzman *et al*. 2010), to more action potentials (Subramaniam *et al*. 2014), or to longer axons with more synapses (Pacelli *et al*. 2015). It has not escaped our notice that faster degeneration in some neurons may be the result of their different palettes of Rab proteins and their effectors.

## Acknowledgements

We are grateful for the gifts of flies from Wanli Smith and the Bloomington Drosophila Supply Center. We also thank Olivia Compton and Martin France who helped with pilot studies, the University of York Biology Technology Facility and Flybase. Ian Martin kindly provided unpublished details of dopaminergic cell loss. We are particularly grateful to Parkinson’s UK and to their volunteers for support (K-1704, G-1804).

## Author contributions

SP, CAM, CU, AF, LC and CJHE performed experiments, CJHE drafted the manuscript, and SP, CAM, CU, AF, LC and CJHE revised the manuscript.

No conflicts of interest were perceived.

